# SISS-Geo: Leveraging Citizen Science to Monitor Wildlife Health Risks in Brazil

**DOI:** 10.1101/286740

**Authors:** Marcia Chame, Helio J. C. Barbosa, Luiz M. R. Gadelha, Douglas A. Augusto, Eduardo Krempser, Livia Abdalla

## Abstract

The well-being of wildlife health involves many challenges, such as monitoring the movement of pathogens; expanding health surveillance beyond humans; collecting data and extracting information to identify and predict risks; integrating specialists from different areas to handle data, species and distinct social and environmental contexts; and, the commitment to bringing relevant information to society. In Brazil, there is still the difficulty of building a mechanism that is not impaired by its large territorial extension and its poorly integrated sectoral policies. The Brazilian Wildlife Health Information System, SISS-Geo, is a platform for collaborative monitoring that intends to overcome the challenges in wildlife health. It aims integration and participation of various segments of society, encompassing: the registration of occurrences by citizen scientists; the reliable diagnosis of pathogens from the laboratory and expert networks; and computational and mathematical challenges in analytical and predictive systems, knowledge extraction, data integration and visualization, and geographic information systems. It has been successfully applied to support decision-making on recent wildlife health events, such as a Yellow Fever epizooty.

## 1 Introduction

Environmental change, including climate change and biodiversity loss, are determining factors for the emergence of diseases originating from wildlife [10] and can be the source of the selective forces of new genetic variations that allow the disruption of biological barriers by pathogens and the increase in the potential for spread of diseases to humans. Although not considered appropriately in health surveillance policies, the situation is relevant, since the majority (60.3%) of infectious diseases circulate between humans and animals (zoonosis), of which 71.8% are caused by pathogens originating from wildlife [18]. Not to mention data from a recent study [37] which makes explicit how delicate the situation is, especially because of the vast and growing number of pathogens infecting humans and animals.

These emergences are widely associated with areas most affected by natural and anthropogenic impacts, also composing the range of parameters that make social inequalities even more severe and unfair, with strong repercussions and costs to health and quality of life [3, 32]. Over the past 15 years several studies have shown the effect of biodiversity in the dilution and dispersion of pathogens and in the modulation of their transmission rate [19, 39, 29].

However, studies and actions in the last century, despite the expansion of epidemiological knowledge, only responded to emergency events of specific diseases in the human population, with some mitigation attempts. Considering the low ability to reverse climate change and environmental impacts determined by human growth, and our way of production and consumption of natural resources, it seems reasonable to expect that we cannot stop the emergence of these diseases. This scenario is paradoxical in megadiverse countries, such as Brazil. While species richness results in richness of parasites that are associated to them, and therefore a potential risk, it is this complexity of species and their relationships that protect and stabilize the dynamics of transmission, reducing the outbreaks of diseases, one of the most important ecosystem services. In this scenario, more than seeking effective responses to crisis situations, there is reason to pursue actions that anticipate problems so that one can mitigate them where possible, and quickly respond to them when prevention or mitigation fail.

This approach has been strengthened with international programs, such as “*One world, one health*” from the WHO/OIE and the 2011-2020 Strategic Plan of the Convention on Biological Diversity (CBD) and strategically in governmental programs of developed countries which already dedicate considerable resources and efforts to tracking pathogens, whether to prevent pandemics, such as the recently occurred with the new influenza and Ebola viruses, the development of new drugs or even biological warfare concerns. In Brazil, systematized strategies for monitoring and predicting occurrences of diseases resulting from biodiversity are incipient. They follow a notification model about diseases that already occurred in humans, which is insufficient for preventive action [5].

The relationships that link biodiversity to health are complex because they are often indirect, scattered in space and time and dependent on numerous forces [29]. The problem is not restricted to identifying species and their geographical distribution. In the context of the emergence of zoonoses there are various species of pathogens, vectors and hosts that modulate evolutionarily each other, their populational dynamics and composition, which collectively also undergo and react to environmental changes [19].

Therefore a multi-dimensional challenge is faced. The first is to sensitize decision-makers about the need to monitor the movement of pathogens in wildlife before they impact humans, expanding health surveillance actions beyond humans. The second dimension is building a mechanism that is not limited by the territorial extension of Brazil, the poorly integrated sectoral policies, and by other outbreaks or emergencies that absorb all the health staff. The third is how to integrate multiple skills, since this mechanism should contain specialists to handle data, species and distinct social and environmental contexts. The fourth is how to effectively obtain information and treat them properly. The fifth is to extract the relevant information from data and to really identify risks and predict them and, finally, the commitment to bring relevant information to society.

As evidenced, data collection, monitoring and extraction of knowledge and information about wildlife health and its relationship to human health arise as challenging tasks involving several areas of knowledge, characterized as interdisciplinary activities aimed at modeling a dynamic and complex system. It is also clear that major areas of computing are essentially applicable in the context presented, such as computer modeling, machine learning and parallel programming; however, their application is not obvious given the need to integrate information in different ways, the complexity and dimensionality of the data to be manipulated and the sensitivity involved in the use and dissemination of these data.

In this article, we present the Information System on Wildlife Health (SISS-Geo), a joint effort between the Oswaldo Cruz Foundation (Fiocruz) and the National Laboratory for Scientific Computing (LNCC), is an important step for moving forward on the challenges posed. Its conception aimed the integration and participation of various segments of society, and encompasses: the registration of primary data by any person interested; the application of the concept of citizen science; the reliable diagnosis of pathogens circulating in wildlife that may potentially impact humans with the participation of laboratory and expert networks; the computational and mathematical challenges that include analytical and predictive systems, data mining, intensive processes, parallel programming, system integration, data (unstructured and heterogeneous) and information, geographic information systems (GIS), machine learning, meta-heuristics, and data visualization.

SISS-Geo is essentially characterized by managing its data in a spatially referenced environment. It aims to:

- provide, quickly and efficiently, the flow of information between (i) the Information Center for Wildlife at Fiocruz and the national system of health surveillance, with special contribution to the Strategic Information Center on Health Surveillance (CIEVS, Ministry of Health); (ii) the participatory networks in wildlife health and laboratories; (iii) the general population that wants to participate in the process; and (iv) the different biodiversity monitoring centers, as the MCTI (Ministry of Science, Technology and Innovation), ICMBio (Chico Mendes Institute for Biodiversity), JBRJ (Botanical Garden of Rio de Janeiro), MAP (Ministry of Agriculture, Livestock and Supply), Embrapa (Brazilian Agricultural Research Corporation), etc.
- create, from the data and georeferenced information, warning and forecasting models on human and wildlife diseases in order to act as a sentinel system for emerging and reemerging diseases as well as provide the results of spatial modeling to scientific community and decision makers.
- allow for adequate means to integrate the georeferenced system with spatial databases partners from governmental and non-governmental partners.
- adapt to the metadata standard of the National Spatial Data Infrastructure (INDE) (http://www.inde.gov.br), aiming to provide, efficiently and with full compatibility, data related to wildlife health to the scientific community and the general population.

## 2 Design and Implementation of SISS-Geo

SISS-Geo is built upon four high-level modules. The first one systematizes the capture of georeferenced field observation records of animals, their physical conditions, and their surrounding environment, which are stored in a database (Sections 2.1 and 2.2). These observations are performed by collaborators through mobile applications, for Android (Figure 1), iOS, and in a web interface (Figure 2). The second module analyzes the data to generate automated alert models that take into account territorial distances, time interval, similarity between taxonomic groups involved (notably for primates, chiroptera, rodents, and carnivores, but not limited to them), the observed physical conditions of the animals in the field according to pre-categorized clinical patterns, and the environmental characteristics of the site where the animal was observed (Section 2.3.1). A georeferenced data explorer is available as well, allowing for multiple layers of information to be overlayed. Figure 3 illustrates a visualization where records (green), alerts (red) and biomes are overlayed in a map of Brazil.

**Figure 1:**
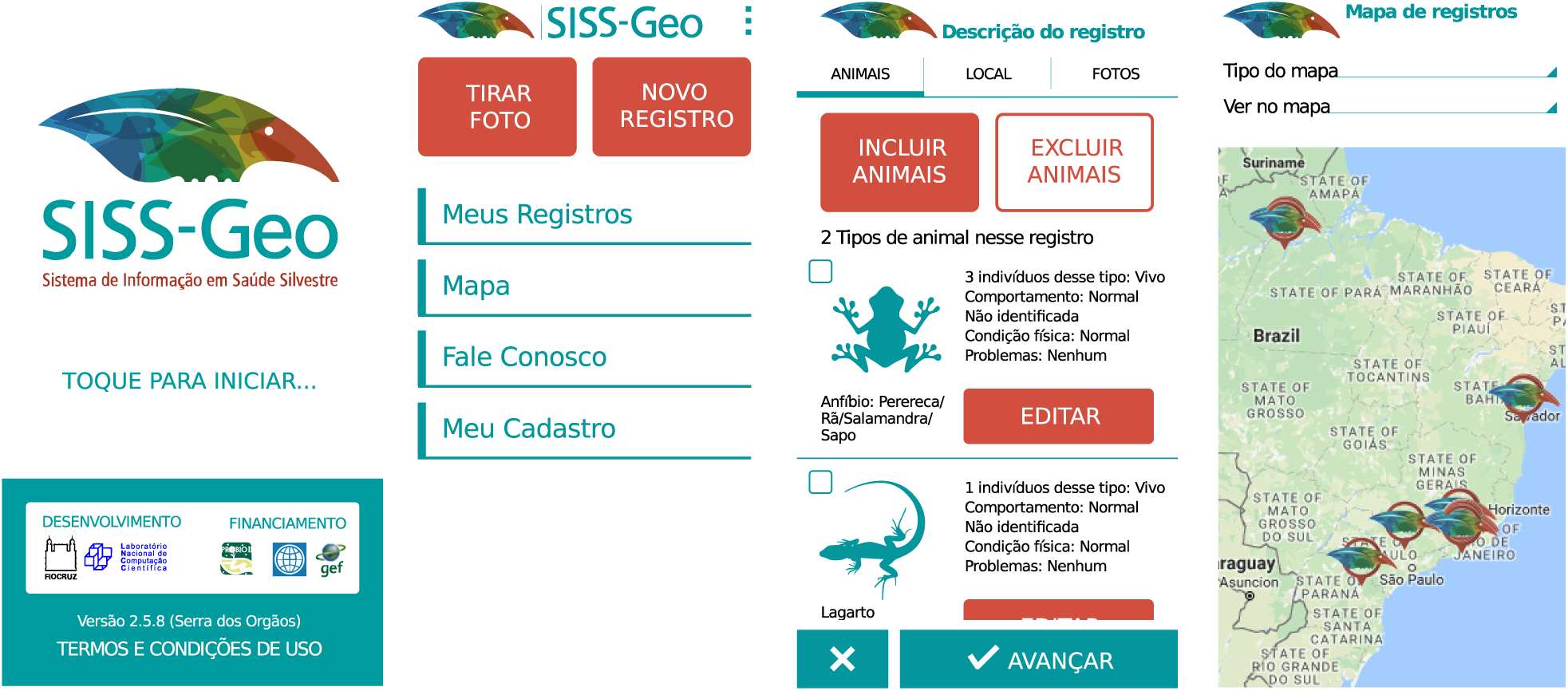
Screenshots of the SISS-Geo mobile application.

**Figure 2:**
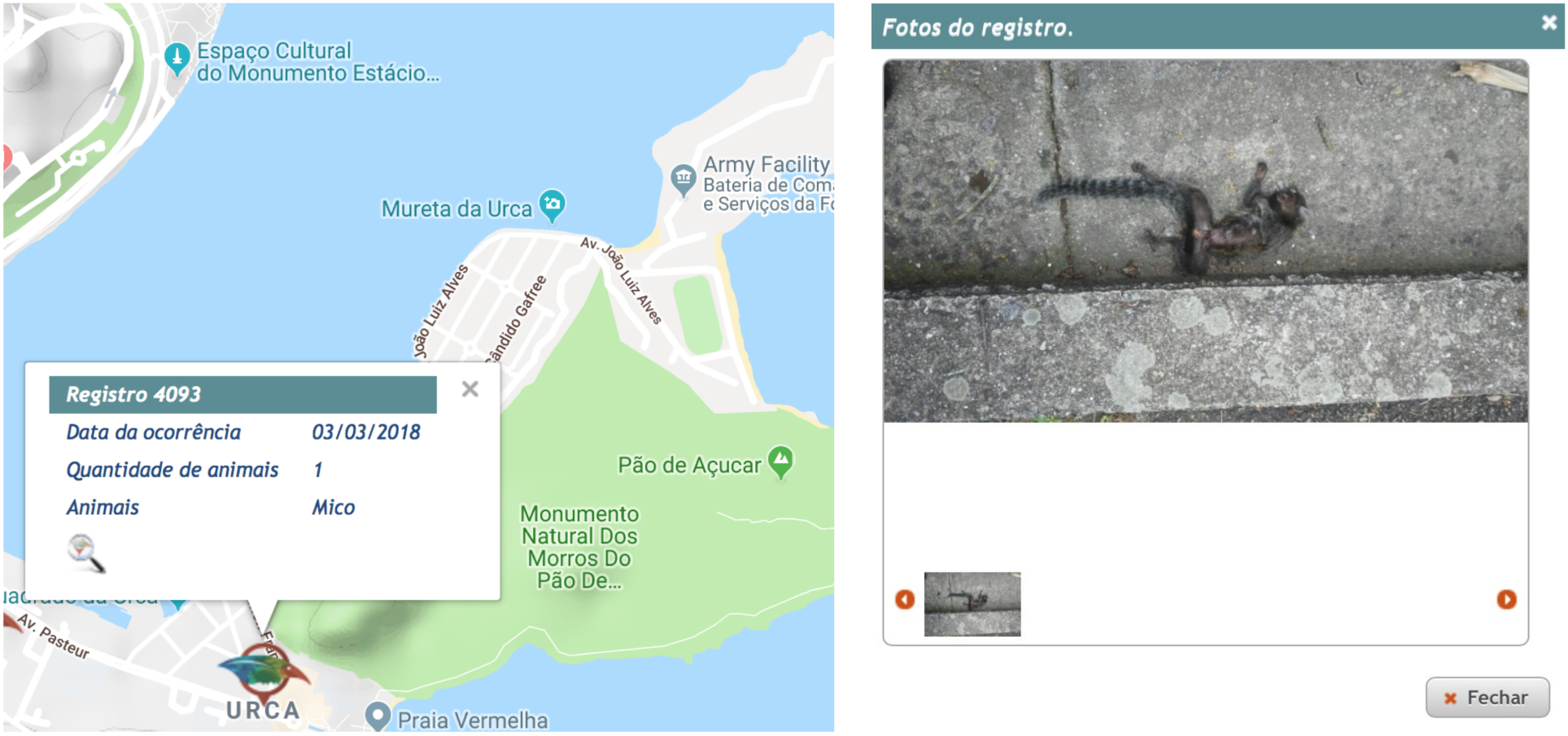
Screenshots of the SISS-Geo web application. Record details in the map (left), corresponding photo with a dead marmoset (right).

**Figure 3:**
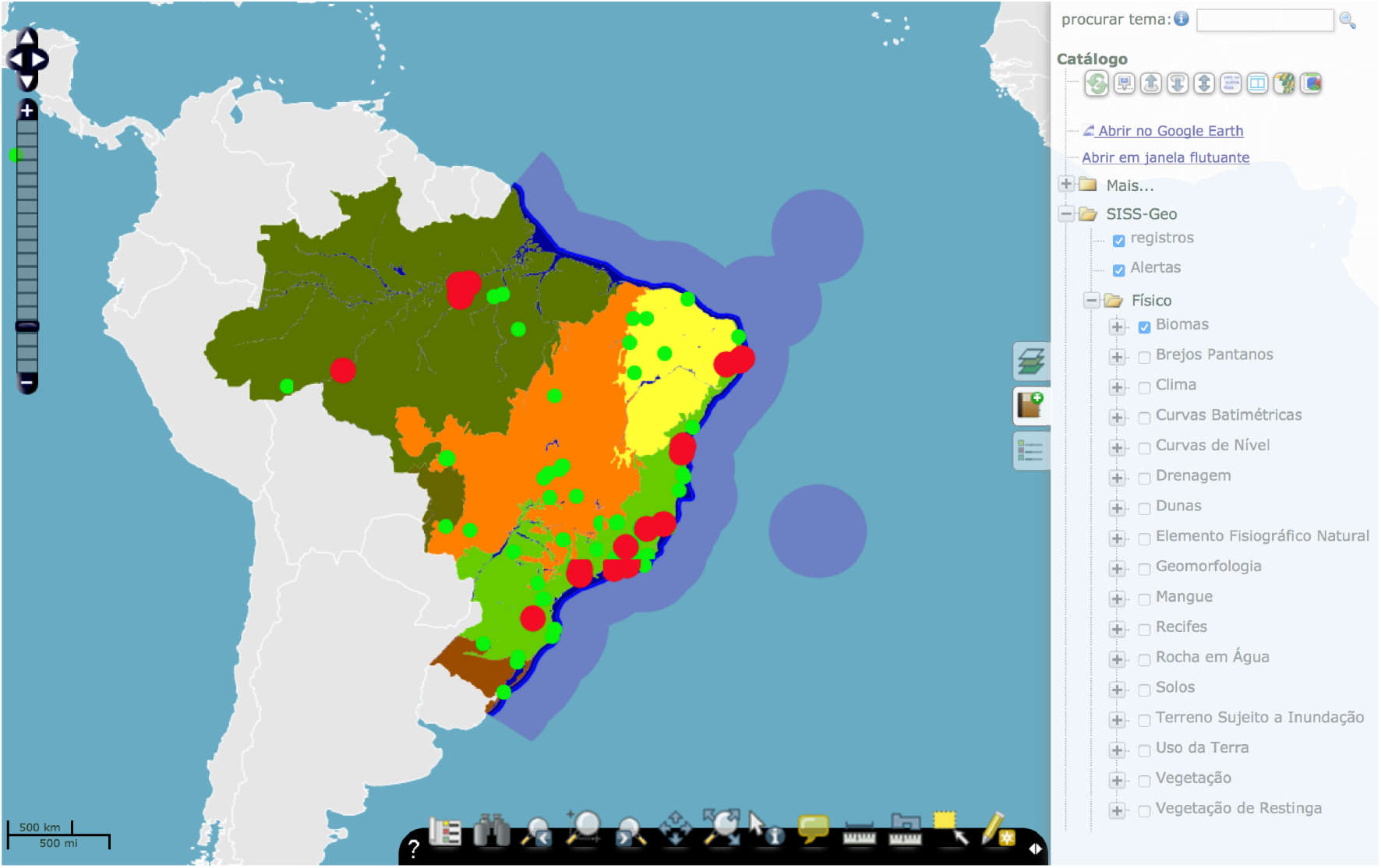
Screenshots of the SISS-Geo georeferenced data explorer.

From the indication of importance and emergency generated by the alert model, the participatory and laboratory networks in wildlife health and the health and environmental services established in the country are requested to collaborate on collecting biological samples in animals in the field and on providing reliable diagnosis. The reliable diagnosis feeds and validates the alert models which in turn, from the initial correlation of the environmental conditions of the occurrence, allows for generation of forecast models of ecological opportunities for disease occurrence that may result from biodiversity loss, thus opening up a different research viewpoint. This is the third module (Section 2.3.2).

Finally, the fourth module approaches the challenge of understanding the relationships that govern the phenomenon in question, from the trained models. In this context, the extraction of knowledge serves as the main hypotheses suggestion mechanism for further investigation and validation by the expert (Section 2.3.3). The main components found in SISS-Geo can be categorized into four classes: wildlife health data management, GIS, machine learning and wildlife health, detailed below.

### 2.1 Data management in wildlife health

To monitor changes in biodiversity, it is essential to collect, document, store and analyze indicators of the spatial and temporal distribution of the species, as well as information on how they interact with each other and with the environment they live in [22]. The development and implementation of mechanisms to produce these indicators [26] depend on access to reliable data from field surveys, automated sensors, biological collections, and from the academic literature. This data is usually available in various institutions that use different formats and identifiers, which makes it a challenging data integration task. The methods and techniques used to manage and analyze this data define a research area often called Biodiversity Informatics [16, 28]. Some initiatives for establishing of metadata and data publishing standards, such as EML [11] and Darwin Core [38], were able to established standard vocabularies used to describe concepts of biodiversity. Although these vocabularies cover only a fraction of possible concepts, they allow institutions to publish their data about biodiversity using the same format, and for their automatic collection and processing by aggregator systems.

Through the use of these standards, SISS-Geo is able to collect species occurrence data provided by various contributors, as well as providing data stored in its own database to the community at large in an easy to use format. Darwin Core has been extended to include concepts on specific topics, such as information about interactions and pollinators (*Darwin Core Extension for Interactions*) and on species profiles (*Plinian Core*). It would be important to evaluate and propose an extension of the standard to include information about wildlife health on species observation records, which is normally carried out in the context of the Biodiversity Information Standards^1^ (TDWG) organization.

SISS-Geo is a biodiversity informatics platform and, as such, it allows for users to upload species occurrence records. In SISS-Geo, these records are enriched with additional attributes, provided by the user, to describe the health condition of the respective individuals. The term *occurrence* is used in this work to refer to the observation of an individual that apparently carries a disease or not, which is a special case of a species occurrence as commonly defined in the biodiversity informatics literature. Its geographical scope is limited to Brazil and the users are given by citizen scientists and specialists. A relational database was conceptually modelled and implemented for SISS-Geo comprising occurrences of organisms along with associated information about their health condition. Standard operations for creating, reading, updating and deleting information are enabled by mobile and web applications that allow for both citizen scientists and system managers to interact with the system (Figure 4 describes SISS-Geo use cases).

**Figure 4:**
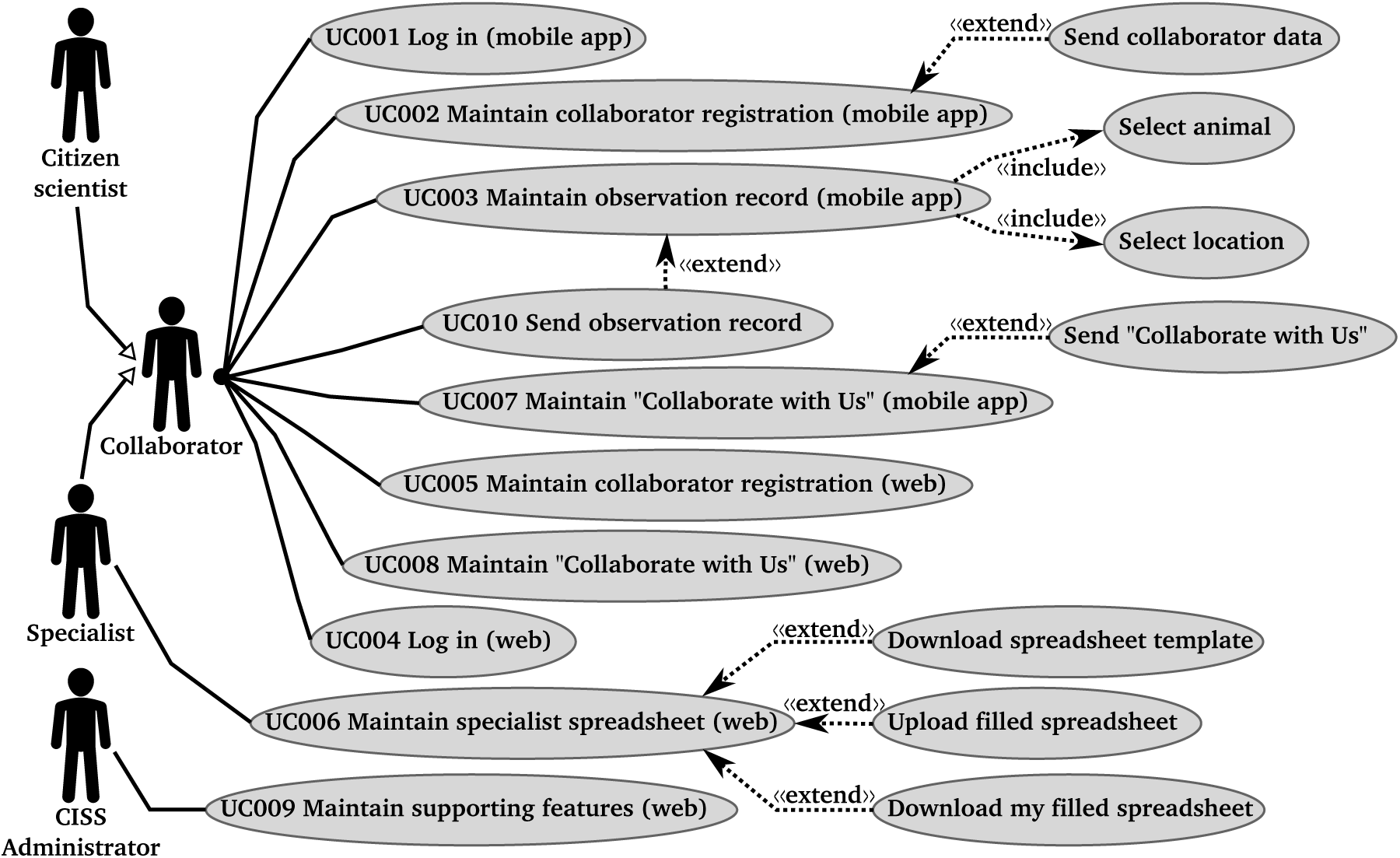
Use cases of SISS-Geo.

As can be observed in its database schema in Figure 5, SISS-Geo stores information about wildlife health occurrences (Occurrence). These occurrences usually have an animal (Animal), a collaborator (Collaborator) and a location (Location) associated to them. Specialists can require samples (Sample) related to the occurrence to be collected, which are be analysed (Analysis) in the laboratory (Laboratory) network. Data stored in this database is consumed by mathematical models that can produce and confirm wildlife health alerts (Alert).

**Figure 5:**
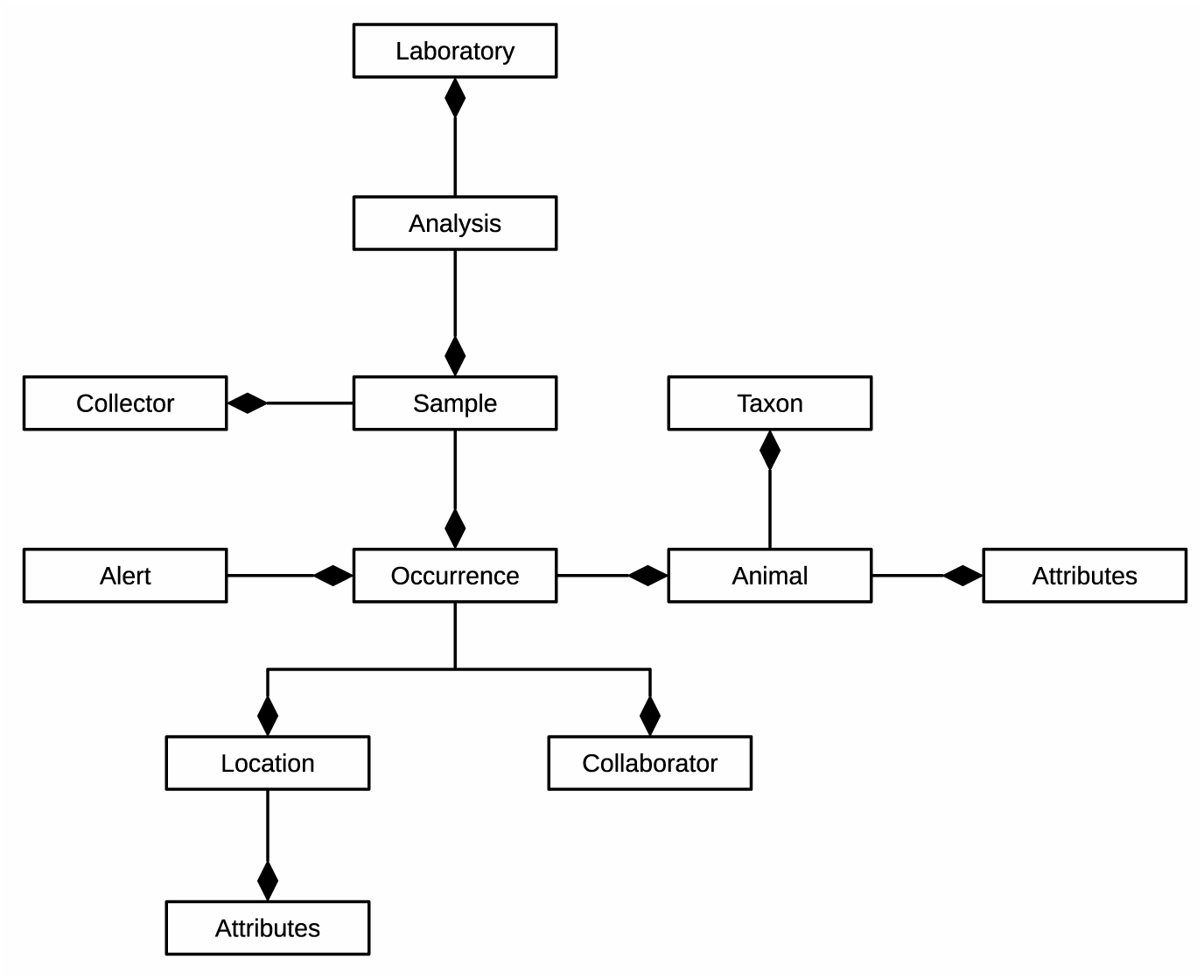
Overview of the database schema of SISS-Geo.

The architecture of SISS-Geo is described in Figure 6. It is comprised by the following components: a mobile application, a web application server, a database server, and high-performance computing resources. As described in the use cases diagram in Figure 4, citizen scientists use the mobile application to request, for instance, the upload of their observations or queries to be executed. These requests are forwarded to the web application server, which connects to the database server to answer these requests. Administrative users and specialists can access the web application server directly to also send requests to SISS-Geo. Finally, the web application server can invoke the execution of computationally-intensive analyzes on high-performance computing resources. A complete list of use cases is described in Figure 4.

**Figure 6:**
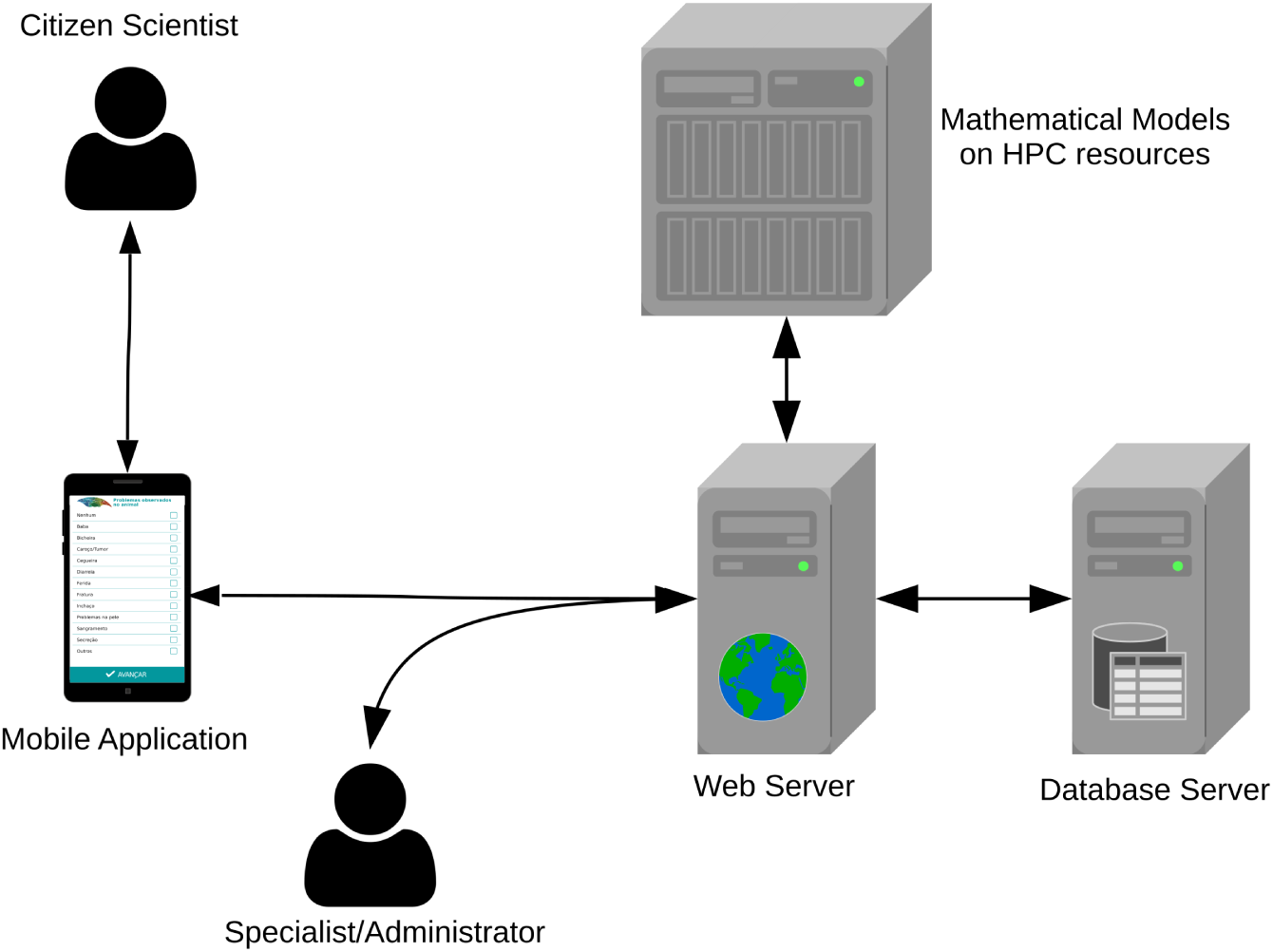
Architecture of SISS-Geo.

### 2.2 Geoprocessing

Spatial and geographical visualization are today basic conditions for the management of information. It is often difficult due to the need for normalization, update and access to qualified data. In studies of infectious diseases, the spatialization of data needs additionally to consider populational pulses and fluctuations determined by several factors such as seasonality, reproductive periods, migrations, among others [25].

SISS-Geo aims to generate relevant and reliable information that is able to support decision processes of the Brazilian Ministries of Health, Agriculture, Livestock and Supply, and Environment, providing subsidies for more agile and timely decision making.

Because it is an innovative project, the functionality developed is not straightforward and it was often not available in similar initiatives. The construction of new methodologies and the use of different types of geographic technologies that can meet the expectations and objectives of SISS-Geo is therefore necessary. The GIS Infrastructure (GI) of SISS-Geo has strategic importance in this process, in which there is a need to overcome challenges related to quality control of spatial data, minimize modeling positional errors, modeling spatialization based on machine learning and the dissemination of models in the form of dynamic maps on the Internet.

The modeling of ecological opportunity of diseases in SISS-Geo uses a big amount of spatial data with different scales, reference systems, sources and mapping methodologies. Therefore it is necessary to normalize and integrate data in a geographic database. This is used both to consume information/data, and to store the modeling results in the form of geographically distributed models. The data used as input for modeling are obtained from the overlap of wildlife occurrence records and environmental, social and human impact databases. Depending on the location of the records, spatial relationships of the types “within”, “close”, “crosses”, etc. can be established.

Systematic databases available from official sources of the Brazilian federal, state and municipal governments, are produced, mostly in small and medium scales (1:1,000,000, 1:500,000). Mapping at these scales provides only a reduced level of detail and accuracy. This can significantly influence modeling in SISS-Geo, since it can result in a degree of uncertainty between the number of points registered and the national cartographic data as they are related.

The uncertainty measurement generally corresponds to the Map Accuracy Standard (MAS)2, whose value is estimated for each map and determines its classification. However, the use of the MAS is questionable when it comes to digital cartography [34], whose development has introduced new mapping techniques and error calculations. The Technical Specification for Vectorial Geospatial Data Acquisition (ET-ADGV), adopted by the Brazilian National Spatial Data Infrastructure (INDE), also addresses this issue and sets new requirements to be followed with respect to the systematic mapping of Brazil. This standard considers that the accuracy in the acquisition of data is equal to the final digital cartographic product because, after vectorial acquisition of an element, its geometry is not modified in subsequent processes. In addition, the accuracy standards considered in this standard are more stringent than those based on analog mapping and are calculated based on statistical comparison of field measurements and the digital product. The adoption of ET-ADGV is a trend, but is still under a process of adaptation, hence few datasets have this information documented. Therefore, the reference of the positional accuracy value of the data for spatial queries of SISS-Geo is initially be based on the MAS.

To minimize the effect of positional error on SISS-Geo’s models takes into consideration the tolerance in space intersections, based on the positional accuracy of the overlay data, using as reference the MAS. This aims to establish models with sufficient positional quality to support decision-making in public health policies.

The geoprocessing infrastructure also needs to make available the results, alerts and prediction models produced by SISS-Geo to the public domain according to the Brazilian Information Access Act, except for sensitive information. Therefore, adequating the geographic information system for the web environment, which provides SISS-Geo’s results in the form of dynamic/interactive maps and graphical statistics, is an ongoing development. An advantage of this technology is the ease of handling, analysis and interpretation of models by the end user, as well as operating system independence and interaction with desktop systems and other Internet systems (interoperability).

### 2.3 Machine Learning

#### 2.3.1 Grouping of observation records and alert prediction

When a wild animal is observed, its physical condition and surrounding environment are recorded in SISS-Geo, either by experts or volunteers. These records are grouped with other related records (previously reported) resulting in a collection of events characterizing a phenomenon. This is the grouping stage and, although it may sound trivial, it involves the challenge of conceiving/training models with discriminative capacity to recognize similarities and dissimilarities between events, based on criteria such as spatial and temporal distance between records, similarity between species and the reported physical conditions, among others. This flow of learning is summarized in Figure 7.

**Figure 7:**
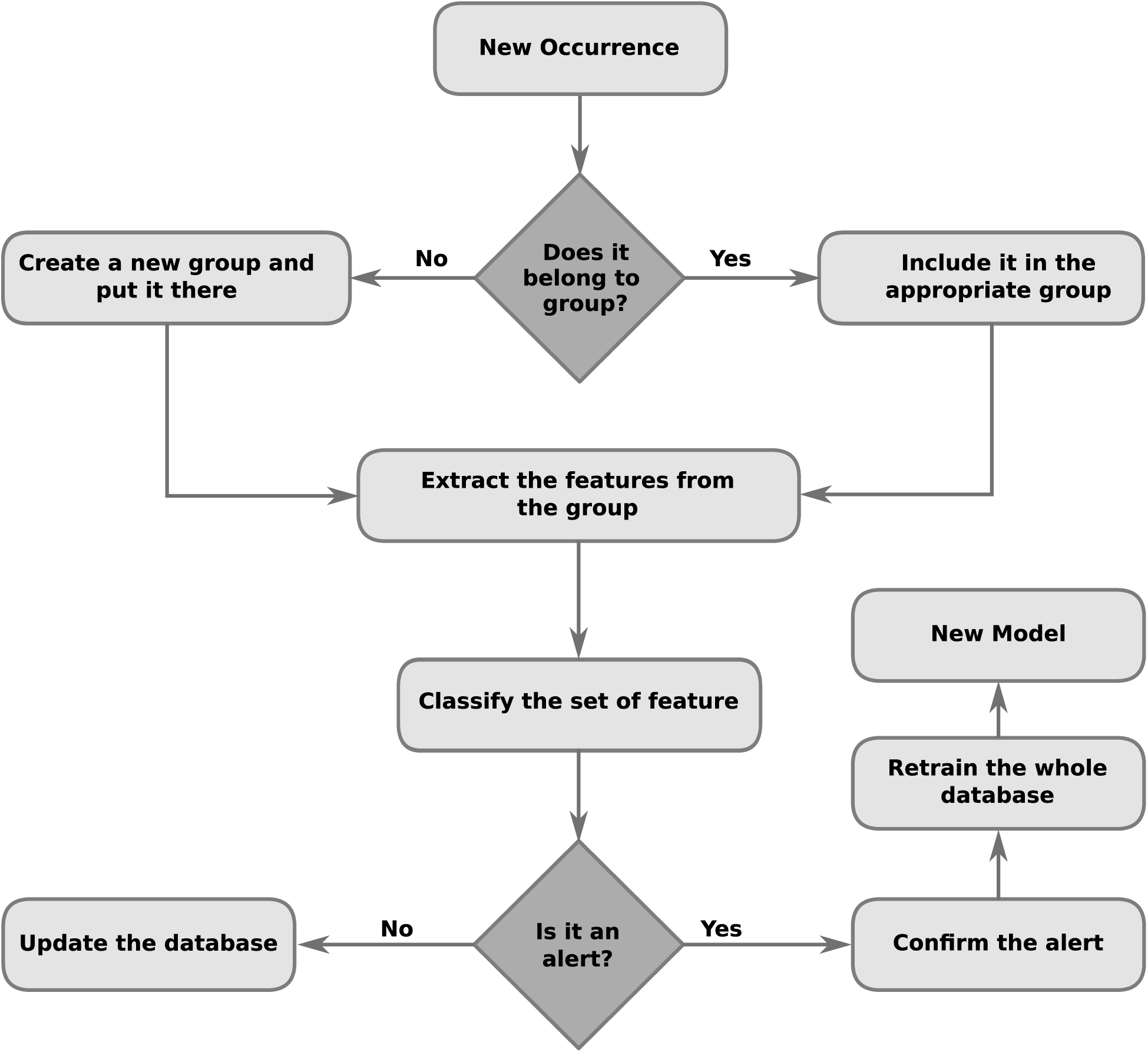
Machine learning flow of SISS-Geo.

The second part consists of modeling the characteristics of observation records that make them more or less relevant, i.e. training the alert model. It means predicting the severity of records according to information brought by events and the geographic/environmental context. For example, a record involving an animal in isolation exhibiting symptoms is less severe, in general, than occurrences containing similar events but covering groups of animals. Of course, in real situations the characterization of an alert situation is usually much less obvious, commonly taking into consideration many factors for decision making. In some cases, a single record is sufficient to generate an alert, such as the registration of a wild canid with symptoms of rabies.

It can be seen that the above mentioned activities refer to the grouping and data classification task, typical of machine learning, and well known for the wide variety of approaches and methodologies. They are therefore complex tasks, both by nature as well as by the large volume of data expected for the system^3^.

However, the challenges of grouping and classification that are present in SISS-Geo go beyond the classic challenges of these tasks.

##### Phenomenon characterization

The characterization of what defines a group of events (phenomenon) lies in the problem of non-conventional similarity measurement formulation (e.g. not necessarily Euclidean). Grouping rules based on expert experience are a reasonable alternative, but has as shortcoming the limited formalization of knowledge and, consequently, the potential for the introduction of unwanted biases. Another approach is to treat this problem as a machine learning process, aiming at the training of similarity models: given a new record and the existing ones, determine to which group it belongs — or whether it characterizes a new group. The process is characterized as supervised learning, since it is possible to determine reliably, *a priori* or *a posteriori*, which records belong to which phenomenon, either by empirical tests or by expert confidence.

##### Feature extraction

Once constituted the phenomena, it is necessary to evaluate them as to the potential threat to wildlife health and its possible outbreak in humans, as phenomena alone do not necessarily constitute alert situations. In this sense, information characterizing a group of events needs to be extracted and provided to the alert prediction model. The difficulty is thus to derive statistics which better represent the phenomenon described by the group in order to maximize the performance of the prediction model; in other words, raise the necessary information to facilitate the learning process. Experts recommend the use of certain statistics, such as the type and quantity of affected animals, number and frequency of occurrences, among others; however, the space of possible features goes well beyond that and could be used to improve predictive performance. Thus, an open question is how to exploit this vast space automatically? An interesting line of research and potential solution to this challenge is the investigation of automatic feature extraction methods [15, 14].

##### Alert prediction model

Although its use in the system is similar to sufficiently known methods described in the literature, the alert prediction model is probably the most strategic component of SISS-Geo’s intelligence. The viability of the system is fundamentally based on the accuracy of the prediction model, both in detecting *true positives* (alerts) as *true negatives* (non-alerts). The failure to detect an alert condition (false negative) can result in serious consequences to wildlife, environmental and human health. On the other hand, false positives would overwhelm the relatively small network of laboratories and experts responsible for confirming or denying alerts (more details below). In this sense, methods that combine multiple models (*ensemble methods*) usually produce more accurate and robust solutions, therefore they are promising candidates as training algorithms for prediction models [31]. Still, since the large portion of the system’s data has no associated class, that is, phenomena whose alert predictions have not yet been confirmed, the semi-supervised learning is an interesting approach due to its ability to also leverage unlabelled instances in the training process [6].

##### Alert confirmation

Another key component of SISS-Geo—in which all others depend—is the process of alert confirmation. A great challenge and bottleneck result from the need for direct human participation in the confirmation procedure, either in the field or laboratory; so it is an expensive and slow process, even considering the extensive network of qualified collaborations linked to SISS-Geo. When there are more alerts issued by the prediction model than the capacity of experts and the laboratory network to confirm them, the phenomena need to be prioritized. In this situation, one can think of prioritizing the phenomena associated with alerts (1) by *alert severity* weighted by *confidence* of prediction; or (2) by *relevance to regions of great interest*, be it social, environmental and/or economic. However, a strategy focused on medium and long term is the prioritization of confirmation (or denial) of alerts with greater potential for improving the accuracy of the prediction model. This line of research is recent and it is called *active learning* [35]. The same method can also be used in possible cases of false negative, thus avoiding the possibility of degeneration of the prediction model^4^: the phenomena predicted as non-alerts but that are promising from the learning point of view would be subject to confirmation (of the non-alert condition) by an expert.

#### 2.3.2 Prediction of Ecological Opportunities for Disease Occurrence

Another line of fundamental importance in SISS-Geo is the prediction of scenarios and environments that favor ecological opportunities for disease occurrence arising from wildlife or, put differently, raising scenarios conducive to the occurrence of a certain event, such as an outbreak of a disease.

In short, trained alert models can be used to evaluate different scenarios and characterize those potentially susceptible. The construction of predictive models should relate various environmental, social, and human and animal health information, being a challenging area to current predictive models. Methods for linking of environmental and animal variables, such as the ones applied to ecological niche modeling or even to more traditional machine learning methods will be widely applied in this context, however, new approaches should be developed, allowing for the integration of the variety of information cited.

Moreover, one should consider the computational challenges involved in the manipulation of information from a large number of records, of different species and environmental conditions, which can be considered a problem with high computational cost. However, despite the expected need to handle large amounts of data, it is also expected for some specific disease or species that only a small number of information will be available, leading to a new challenge: applying prediction techniques in a highly unbalanced setting.

#### 2.3.3 Knowledge Extraction

An important feature of symbolic modeling methods, such as decision trees, rule extraction algorithms and meta-heuristic genetic programming [20], is that the model is itself the explicit representation of knowledge extracted from data. More specifically, it is the revelation—subject to human interpretation—of the existing relationships between the input and output data.

It is remarkable the potential of this class of models to aid experts in the analysis and understanding of the phenomenon investigated, leading to a man-machine interaction: the model suggests hypotheses that best fit the data while the expert validates them.

The central challenge of knowledge extraction is in defining the structure/language of the model or, in other words, the incorporation of expert knowledge. Therefore, it is also challenging to find the ideal balance between *bias*, usually resulting from structural simplicity of the model, and *variance*, an issue usually associated with structurally more complex models.

Depending on the parametrization of the learning algorithm and dimensionality of the database, a second challenge arises: the associated computational demand is exacerbated by the fact that the process of knowledge extraction is often performed interactively by the specialist. Typically, the strategies employed in these situations include (1) reducing the dimensionality of the data [12] and (2) leveraging parallel and distributed computing, in conventional architectures or accelerators [1].

## 3 Evaluation

As of February 2018, SISS-Geo was downloaded more that a thousand times from the Google Play store and had an average rating of 4.8 out of 5 starts. Even though the potential number of observations related to wildlife health usually being a fraction of the population of a species, SISS-Geo has 3,014 records in its database performed by 1,881 citizen scientists. Its web interface has been accessed 4,463 times. These records correpond to 764 mammals, 815 birds, 383 reptiles, 227 amphibians, 47 fish, and 540 not identified. Table 1 lists the ten most recorded common names in SISS-Geo. It is important to emphasize that the records were uploaded by volunteer collaborators that often do not have taxonomic knowledge, which can have adverse effects on data quality. To tackle this issue and improve wild animal monitoring, which can lead to better assertive models for the emergence of zoonoses, SISS-Geo has developed a tool for expert-supported record validation. Figure 8 shows the geographic distribution of the observations recorded by SISS-Geo that are georeferenced.

**Table 1:** Ten most recorded common names in SISS-Geo

| Common name | Number of records | % |
| --- | --- | --- |
| Not identified | 540 | 18.3 |
| Birds (Other) | 417 | 14.1 |
| Birds (Penelopes, Seriemas, Toucans) | 112 | 8.6 |
| Amphibians | 242 | 8.2 |
| Snakes | 189 | 6.4 |
| Marmosets and Tamarins | 177 | 6.0 |
| Turtles | 101 | 5.9 |
| Lizards | 75 | 3.4 |
| Birds of prey | 72 | 2.5 |
| Capybaras | 64 | 2.2 |

**Figure 8:**
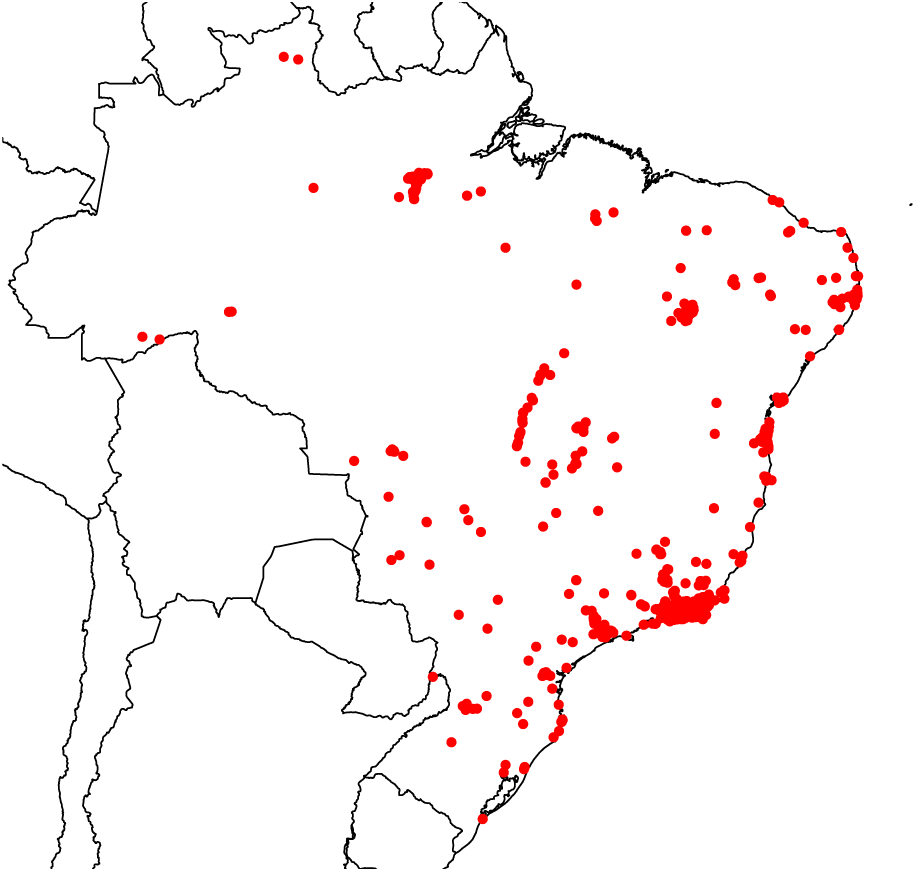
Geographic distribution of records (red dots) in Brazil.

SISS-Geo integrates data-based computational modeling, development and high-performance computing. It was selected in 2014 as the best project [4] in the “Health” category of the *Grand Challenges of Computing* event of the Brazilian Computer Society. In 2017, SISS-Geo received the National Biodiversity Prize from the Brazilian Ministry for the Environment^5^. It allows the monitoring of wildlife and can support the identification of zoonoses, such as the Yellow Fever zoonosis, which in its wild cycle circulates among primates. The fact that monkeys become ill or die before there are human cases of Yellow Fever causes the surveillance of outbreaks, such as the recent one [8, 24], in these animals to be of major importance in the control and prevention of the disease. The collaboration of the population is very important, because prevention actions can be improved and streamlined and everyone will benefit. With the participation of ordinary people, the application makes available, in real time, the occurrences of dead or diseased animals for public health and biodiversity conservation, assisting the Epizootics Surveillance System in Nonhuman Primates (PNH), of the Brazilian Ministry of Health, and records of dead monkeys are reported to the responsible bodies investigating the cases. The information recorded in SISS-Geo serves to generate computational models for predicting zoonoses and for the adoption of preventive measures. Tables 2 and 3 list the recorded conditions and the most recorded abnormalities in SISS-Geo respectively.

**Table 2:** Recorded conditions in SISS-Geo

| Condition | Number of records | % |
| --- | --- | --- |
| Normal behavior | 2,168 | 71.2 |
| Dead animal | 697 | 22.9 |
| Strange behavior | 112 | 3.7 |
| Sick animal | 67 | 2.2 |

**Table 3:** Recorded abnormalities in SISS-Geo

| Abnormality | Number of records | % |
| --- | --- | --- |
| None | 2,377 | 81.3 |
| Wound | 103 | 3.5 |
| Other | 101 | 3.5 |
| Fracture | 75 | 2.6 |
| Blindness | 74 | 2.5 |
| Bleed | 58 | 2.0 |
| Skin problems | 36 | 1.2 |
| Swell | 32 | 1.1 |
| Myiasis | 25 | 0.9 |
| Secretion | 18 | 0.6 |
| Drool | 16 | 0.5 |
| Lump or Tumor | 6 | 0.2 |
| Diarrhea | 3 | 0.1 |

Some of the observations performed with SISS-Geo triggered alerts and contributed to biodiversity conservation actions, such as: (i) 59 dead turtles were recorded in the south of the Brazilian state of Bahia, generating a notification to the responsible environmental agency and a legal notice to those involved in predatory fishing in the area; (ii) 73 dead monkeys were recorded during the recent Yellow Fever epizooty, which directed health surveillance actions in the field; (iii) observations of dead foxes with rabies in the Northeast were able to support decision-making by health surveillance agencies.

SISS-Geo also contributed to the monitoring of species on the IUCN Red List of Threatened Species, with the availability of the location and information of some species already registered as: *Panthera onca, Puma concolor, Tapirus terrestris, Myrmecophaga tridactyla, Bradypus torquatus, Chrysocyon brachyurus* e *Chelonia mydas*; *Leontopithecus chrysomelas, Alouatta guariba guariba, Crax blumenbachii*.

## 4 Related Work

He et al. [17] present the eMammal framework for wildlife monitoring supported by citizen scientists. Animal images collected with camera traps are sent to its database where visual animal recognition techniques are applied. The species identification recommendations generated are reviewed by citizen scientists and, subsequently, by experts. The resulting validated records are made available to wildlife and ecological researchers. eBird [36] also leverages the capability of citizen scientists to gather bird observation records. Automated data quality filters are used to support species identifications performed by citizen scientists.

More general biodiversity databases exist at the global, national and ecosystem levels. GBIF [9] gathers species observation data at a global scale. In February 2018, it had 54 national nodes. Along with other types of participants, GBIF gathers data from 1.152 institutions, totaling approximately a billion records. SiBBr [13] is the Brazilian GBIF node, publishing species occurrence records and providing an ecological niche modeling portal [33]. BaMBa [21] is a biodiversity database that focuses on marine ecosystems that is also integrated with GBIF. These systems use the IPT tool [30] to extract observation records from local databases, export them to Darwin Core [38], and publish them on GBIF.

SISS-Geo is both a citizen science application and a biodiversity database. eBird and eMammal, while being citizen science applications as well, do not provide tools for data analysis as SISS-Geo does with the application of machine learning techniques to generate wildlife health alerts. GBIF, SiBBr, and BaMBa focus on data mobilization and publication and do not directly provide tools for enabling the participation of citizen scientists.

## 5 Conclusion

The proposal was inspired by the desire to make public and seek reinforcements for a long walk that brings together researchers, experts from multiple areas and society so that, through computing, information and disease prevention actions reach the most remote regions of the country. It emerges from many years of practice of field research in the Brazilian semi-arid region, where relevant information on diseases in wild animals have been lost or dispersed and the lack of systematization turned important actions impossible both for the containment of diseases in humans, as for conservation of species.

SISS-Geo was born of efforts to create innovative and integrated actions for the mainstreaming of biodiversity in the sectors of the country. It integrates the actions of the Oswaldo Cruz Foundation (Fiocruz) in “Public-Private Actions for Biodiversity Project” – PROBIO II^6^, coordinated by the Brazilian Ministry of Environment, and developed by FUNBIO, Embrapa, the Brazilian Ministries of Agriculture and Livestock, Health, and Science Technology and Innovation, the Botanical Garden of Rio de Janeiro, ICMBio and Fiocruz. The National Laboratory for Scientific Computing joined the Fiocruz project and ensured its execution in a long-term knowledge-building partnership.

By automating the search for occurrence patterns, the information reaches more efficiently citizens nationwide, from the general population through experts, as well as provides the opportunity for the acquisition of knowledge about the possible patterns and parameters that contribute to the occurrence of diseases. In the medium- and long-term it also builds the capacity of researchers to develop complex modeling in ecology of diseases that can possibly exploit geographic information in order to improve accuracy. Moreover, occurrence patterns yield data that can assist national policy on health and on biodiversity conservation.

In the context of SISS-Geo, we plan to incorporate provenance information to allow the alert generation process to be traceable, meaning that one can recover the data, configuration parameters, people and computational activities involved. This enables many applications, such as assessing the quality of the alerts generated, verifying compliance with governmental regulations, and the reproducibility [7, 2] of the alert generation process. Provenance information [23, 27], which contains details about the planning and execution of computational processes, such as scientific workflows, describing the processes and data involved in the generation of its results may be used to facilitate this task. They allow precise description of how a computational process was planned, which is called *prospective provenance*, and what occurred during execution, which is called *retrospective provenance*. Some applications of provenance include reproducibility of computational processes for validation, sharing and reuse of knowledge, data quality evaluation and attribution of scientific results. One of the concepts commonly captured in provenance is *causality*, which is given by the existing dependency relationships between computational activities and data sets. These dependencies can derive, by transitivity, dependencies between data sets and between processes.

## Availability

Web interface:

http://sissgeo.lncc.br

Geographical explorer:

http://morcego.siss.lncc.br/i3geo/interface/black_ol.htm

Mobile application (Android): https://play.google.com/store/apps/details?id=siss.ui

Mobile application (iOS): https://itunes.apple.com/br/app/siss-geo/id1291912325

## Footnotes

1 http://www.tdwg.org

2 In Portuguese “Padrão de Exatidão Cartográfica (PEC)”; it is a Decree-Law regulatory technical standards of Brazilian mapping.

3 After all, it is an ambitious system that aims to aggregate and store records on wildlife health of a vast country.

4 Consider the extreme situation where all the predictions are *non-alerts*, including both true as false negative. Since in principle only the cases of alerts are of interest and subject to confirmation, in this scenario the model would be doomed to degeneration.

5 http://www.mma.gov.br/index.php/comunicacao/agencia-informma?view=blog&id=2349%20 (In Portuguese)

6 http://www.funbio.org.br/probioii

